# MAGpy: a reproducible pipeline for the downstream analysis of metagenome-assembled genomes (MAGs)

**DOI:** 10.1101/233544

**Authors:** Robert Stewart, Marc Auffret, Tim Snelling, Rainer Roehe, Mick Watson

## Abstract

Recent advances in bioinformatics have enabled the rapid assembly of genomes from metagenomes (MAGs), and there is a need for reproducible pipelines that can annotate and characterise thousands of genomes simultaneously. Here we present MAGpy, a Snakemake pipeline that takes FASTA input and compares MAGs to several public databases, checks quality, assigns a taxonomy and draws a phylogenetic tree.

## Introduction

Discovering and studying microbes in the environment has been a goal of genomic technologies for many years [1,2], but advances in DNA sequencing [3–5] have enabled a revolution in metagenomics that has accelerated this area of research. Metagenomics refers to the whole-genome investigation of every organism within a particular environment, and is often used in microbiome studies to investigate changes in the taxonomic and functional profile of samples of interest. This method of simultaneously quantifying taxonomic and functional structure has been used in studies of age and geography in the human gut [6], release of carbon due to permafrost thawing [7], the environmental impact and feed efficiency of animal agriculture [8,9], environmental characterisation of Earth’s oceans [10,11] and the extraction of industrially and commercially relevant enzymes from environmental samples [12].

Metagenomics also offers the ability to assemble near-complete and draft microbial genomes without the need for culture. Such “metagenomic binning” approaches involve the assembly of metagenomic sequence reads into contigs followed by clustering, or binning, of contigs into putative genomes, called metagenome-assembled genomes (MAGs) [13]. This was first demonstrated in 2004 when Tyson *et al* [14] assembled multiple near complete genomes from an environmental biofilm. More recently, this technique has been used to assemble complete and draft genomes from ruminants [15–18], oil reservoirs [19], aquifers [20] and the Tara oceans expedition [21].

A major challenge in the analysis of metagenome-assembled genomes is that researchers are often presented with hundreds or thousands of putative genomes, which need to be annotated, characterised, placed within a phylogenetic tree and assigned a putative function or role. This is further complicated by the fact that many of the putative genomes do not have close relatives with good quality reference genomes, making comparative genomics almost impossible.

Here we present MAGpy (pronounced ‘magpie’), a reproducible pipeline for the characterisation of MAGs using open source and freely available bioinformatics software. MAGpy is implemented as a Snakemake [22] pipeline, enabling reproducible analyses, extensibility, HPC integration with high-performance-compute clusters (HPC) and restart capabilities. MAGpy annotates the genomes, predicts putative protein sequences, compares the MAGs to multiple genomic, proteomic and protein family databases, produces several reports and draws a taxonomic tree. We demonstrate the utility of MAGpy on a subset of 800 bacterial and archaeal MAGs recently published by Parks *et al* [23].

## Methods and results

Snakemake is a workflow management system with a python-based workflow definition language, and works with any tool, including web services, that have well defined inputs and outputs. Workflows are defined within Snakefiles, within which “rules” are defined that are essentially operations on data, converting inputs to outputs. That conversion can be carried out by a locally installed tool, or shell or Python code embedded within the Snakefile itself. The workflow and rule dependency is defined by setting the output of one tool as the input to another. Snakemake scales to 1000s of input datasets and complex workflows as it integrates with many HPC job management platforms, such as SGE or Slurm.

MAGpy makes use of Snakemake to define an analysis workflow for MAGs based on open source and freely available bioinformatics software. First, CheckM [24] is run, which uses a set of pre-computed core genes to assess the completeness and contamination of MAGs. CheckM also attempts to assign a taxonomic level to the MAGs, though in our experience this is often a conservative estimate. In tandem, MAGpy predicts the protein coding sequences of MAGs using Prodigal [25]. DIAMOND [26] blastp is used to compare the proteins to UniProt [27]. This has multiple purposes – the hits from UniProt provide a form of annotation of the putative proteins and may predict function; many MAGs may show little similarity to published genomes at the DNA level, but as proteins are more conserved, protein hits can help define the closest sequenced genome; and the length of the predicted protein and that protein’s hits can be used to detect truncated genes and proteins in the MAG annotation. Reports of the DIAMOND results at the level of the MAG and for each contig within each MAG are produced. The proteins are also compared to protein families in Pfam [28] using PfamScan; and to draw a tree using PhyloPhlAn [29], which is subsequently visualised using GraPhlAn [30]. The MAG genome sequences are also compared to over 100,000 public genomes using MinHash signatures as implemented in Sourmash.

We applied MAGpy to 800 Bacterial and Archaeal MAGs from Parks *et al* [23]. The CheckM report (which uses Ete3 [31] to expand the taxonomic prediction) can be seen in Supplementary Table 1, the Sourmash report in Supplementary Table 2 and the Uniprot report in Supplementary Table 3. The PhyloPhlAn tree can be seen in Figure 1.

**Figure 1.**
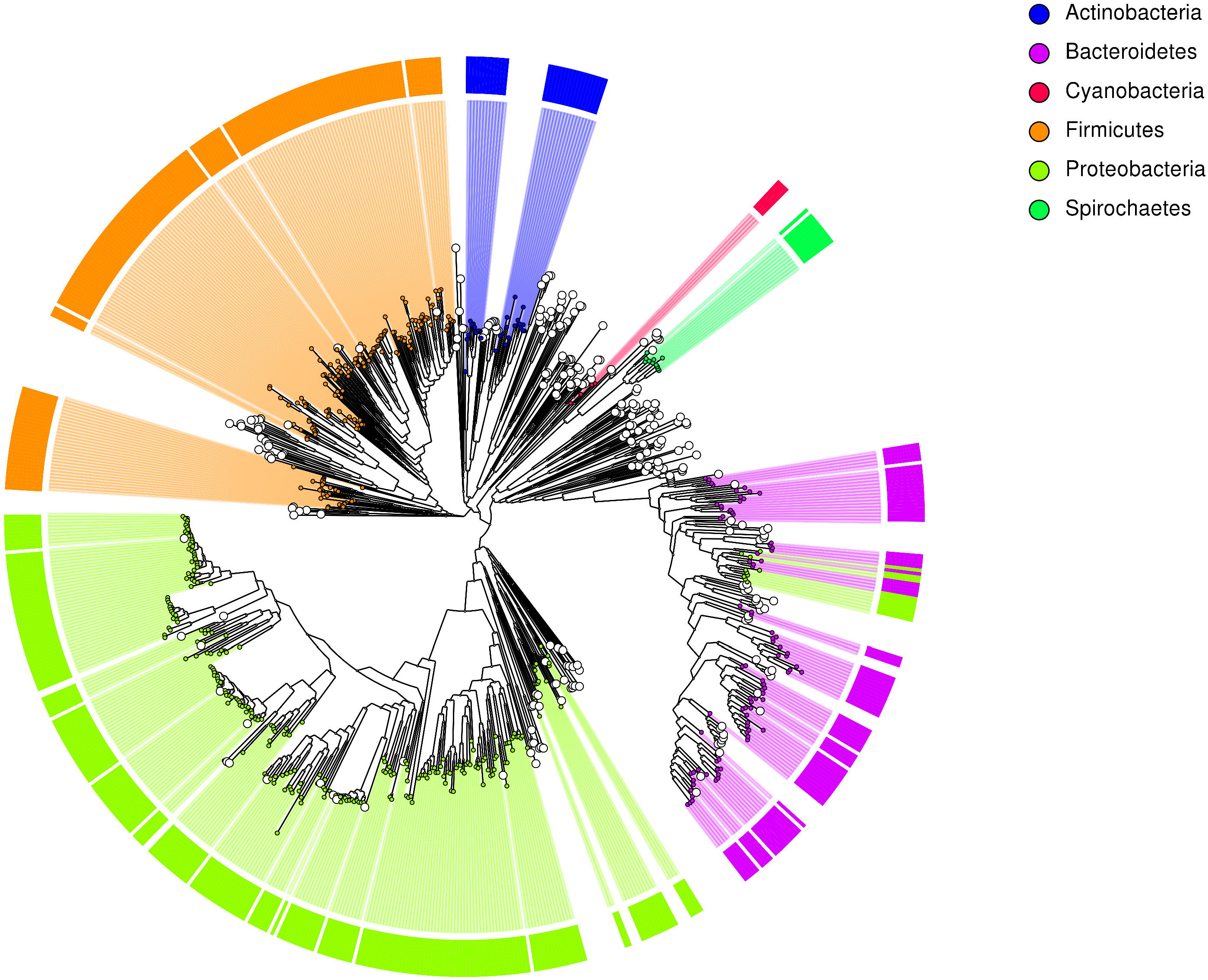
Phylogenetic tree of 800 MAGs created using PhyloPhlAn and produced by MAGpy.

## Discussion and conclusion

Given the cost and difficulty of microbial culture, being able to assemble whole genomes from metagenomes without the need for culture is a hugely promising method which will enable researchers to populate the microbial tree of life rapidly and at relatively low cost. Short read assembly and binning are currently the most powerful method for creating MAGs; however, as long reads become more readily used for environmental metagenomes [32], this will enable the assembly of complex and difficult to assemble genomes directly from the environment, including those with large numbers of repeats and rearrangements [33,34]. Whichever sequencing technology is used, we predict that a huge amount of novel microbial genomes will need to be annotated and analyzed in a consistent and reproducible way, and publication of such pipelines is a key aspect of open science [35].

## Availability

All code and documentation is available at <u>https://github.com/WatsonLab/MAGpy</u>

## Acknowledgements and funding

The Roslin Institute forms part of the Royal (Dick) School of Veterinary Studies, University of Edinburgh. This project was supported by the Biotechnology and Biological Sciences Research Council (BBSRC; BB/N016742/1, BB/N01720X/1), including institute strategic programme and national capability awards to The Roslin Institute (BBSRC: BB/P013759/1, BB/P013732/1, BB/J004235/1, BB/J004243/1); and by the Scottish Government (RESAS) as part of the 2016-2021 commission.

